# The rice G protein γ subunit *qPE9-1* positively regulates grain-filling process by interacting with abscisic acid and auxin

**DOI:** 10.1101/487850

**Authors:** Dongping Zhang, Minyan Zhang, Yong Zhou, Yuzhu Wang, Hongyingxue Chen, Weifeng Xu, Lin Zhang, Ting Pan, Bing Lv, Guohua Liang, Jiansheng Liang

**Affiliations:** Jiangsu Key Laboratory of Crop Genetics and Physiology/Co-Innovation Center for Modern Production Technology of Grain Crops, Key Laboratory of Plant Functional Genomics of the Ministry of Education, Yangzhou University, Yangzhou, China, 225009; Department of Biology, Southern University of Science and Technology, Shenzhen, China 518055; College of Life Sciences and Key Laboratory of Ministry of Education for Genetic Breeding and Multiple Utilization of Crops, Fujian Agriculture and Forestry University, Fuzhou, Fujian, China 350002

**Keywords:** G-protein, *qPE9-1*, starch biosynthesis, grain filling, hormones, *Oryza sativa*

## Abstract

The rice genome contains a single G_α_ (*RGA1*) and G_β_ (*RGB1*) and five G_γ_ subunits. Recent genetic studies have shown that *DEP1/qPE9-1*, an atypical putative G_γ_ protein, is responsible for dense and erect panicles, but the biochemical and molecular mechanisms underlying control of grain size are not well understood. Here, we report that plants carrying *qPE9-1* have more endosperm cells per grain than plants contain the *qpe9-1* allele. The *qPE9-1* line has a higher rate and longer period of starch accumulation than the *qpe9-1* line. Additionally, the expression of several key genes encoding enzymes catalyzing sucrose metabolism and starch biosynthesis is higher in the *qPE9-1* line than in the *qpe9-1* line, especially from the mid to late grain-filling stage. Grains of the *qPE9-1* line also have higher contents of two phytohormones, ABA and IAA. Exogenous application of ABA or IAA enhanced starch accumulation and the expression of genes encoding grain-filling-related enzymes in the grains of *qPE9-1*, whereas only IAA produced these effects in *qpe9-1*. Based on these results, we conclude that *qPE9-1* promotes endosperm cell proliferation and positively regulates starch accumulation largely through ABA and IAA, which enhance the expression of genes encoding starch biosynthesis during the late grain-filling stage.

## Introduction

Rice is the major staple food for more than half of the global population. This crop provides approximately 20% of the global dietary energy supply and plays vital roles in sustaining global food security. Understandably, improving rice productivity is the predominant target in rice breeding because of the continuing increase in the global population. Rice yield is a complex trait that is influenced by numerous components, including the number of panicles per plant, the number of grains per panicle and the grain size (i.e., 1000-grain weight). Furthermore, these components interact with each other in most cases. Typically, the rice grain yield is determined by both the grain capacity (i.e., the grain number and size) and the grain-filling efficiency; the former is controlled by the numbers of spikelets per panicle and the numbers and sizes of endosperm cells per spikelet/grain, both of which determine the final grain number per panicle and the grain size. The grain-filling efficiency is closely related to the grain-filling rate and duration.

The heterotrimeric G protein (hereafter G protein)-mediated signal transduction pathway is considered one of the most important signaling mechanisms and regulates various important physiological and molecular processes in both mammals and higher plants (Urano *et al.*, 2014). In this signaling pathway, the G protein, which is well-known to consist of three different subunits (α, β and γ) acts as a signal intermediator in the transduction of numerous external signals (Milligan & Kostenis, 2006). Arabidopsis G-protein Gamma subunit 3 (*AGG3*), which represents a novel class of canonical γ subunits in Arabidopsis, is widely spread throughout the plant kingdom but is not present in animals (Urano *et al.*, 2013). Recently, *AGG3* has been proposed to be an important regulator of organ size and a mediator of stress responses in *Arabidopsis*. Roy Choudhury *et al.* (2013) overexpressed *AGG3* in *Camelina* and found that it increased the seed size, seed mass and seed number per plant by 15%-40%, effectively resulting in a significantly higher oil yield per plant. In addition, recently *AGG3* has been shown to affect the guard cell K^+^ channel, morphological development, ABA responses and cell proliferation (Chakravorty *et al.*, 2011; Li *et al.*, 2012; Roy Choudhury *et al.*, 2013). These observations draw a strong link between the roles of *AGG3* in regulating two critical yield parameters (seed traits and plant stress responses) and reveal an effective biotechnological tool to dramatically increase agricultural crop yield.

Homologs of *AGG3* in rice have been identified as important quantitative trait loci (QTL) for grain size and yield. The major rice QTL [i.e., *GRAIN SIZE 3* (*GS3*) and *DENSE AND ERECTPANICLE1* (*DEP1*)/*PANICLE ERECTNESS* (*qPE9-1*)] have 29.4% and 22.5% amino acid sequence identities, respectively, with *AGG3* (Li *et al.*, 2012). As an atypical G_γ_ protein, rice *GS3* has been proposed to negatively affect cell proliferation, whereas *DEP1/qPE9-1* plays a positive role (Fan *et al.*, 2006; Huang *et al.*, 2009). These findings suggest that *AGG3* and its homologs in rice may have divergent functions.

In the present study, we show that *qPE9-1* stimulates endosperm cell proliferation, grain filling and consequently the final grain weight. The results also indicate that *qPE9-1* exerts its effects at least in part by controlling abscisic acid (ABA) and indole-3-acetic acid (IAA) biosynthesis.

## Materials and Methods

### Plant material and treatments

The experiment was carried out at the farm of Yangzhou University (32°30’N, 119°25’E) during the rice (*Oryza sativa*) growing season (from early May to early September). Transgenic rice *qPE9-1* and the donor Wuyunjing 8 (*qpe9-1*) were grown in the field (see Zhou *et al.*, 2009). At the heading stage, 500 uniformly growing and headed panicles (1-2 panicles per plant) were chosen, and spikelets on the selected panicles with the same flowering date were labeled for each cultivar/line. The flowering date and position of each spikelet on the labeled panicles were recorded. Approximately 45 labeled panicles were sampled at each time point from flowering to maturity. Half of the sampled grains were frozen in liquid nitrogen for at least 2 min and then stored at −80°C for subsequent analyses. The other half of the grains were dried at 80°C for approximately 72 h to a constant weight and used for the starch analyses.

The hormone treatment consisted of 80 plastic pots with planted rice (three hills per pot) maintained under open field conditions. Each pot (0.6 m in height with 0.5 m and 0.3 m top and bottom diameters, respectively) was filled with sandy loam soil that contained the same nutrient contents as the field soil. The sowing date and cultivation were the same as those for the field experiment. After flowering, either 50 μM ABA or 110 μM IAA was sprayed at a rate of 10 ml per pot on the top of the plants (spikes) every 3 days. Both ABA and IAA were applied between 16:00 h and 18:00 h. All of the solutions contained ethanol and Tween 20 at final concentrations of 0.1% (v/v) and 0.01% (v/v), respectively. The control plants were sprayed with the same volume of deionized water containing the same ethanol and Tween 20 concentrations. Each treatment consisted of 13 pots, and the labeled spikelets were sampled.

### Isolation and counting of endosperm cells

The endosperm cells of the grains were isolated and counted according to the procedures described by Singh & Jenner (1982). Briefly, fixed grains (10 grains from five labeled panicles) were transferred to 0.7:1 (v:v) ethanol:water and dehulled. The dehulled grains were transferred to 0.5:1 (v:v) ethanol, 0.25:1 (v:v) ethanol and finally to distilled water for 5~7 h prior to dissection of the endosperms. The endosperms were isolated under a dissecting microscope and dyed using Delafield’s hematoxylin solution for 24~30 h, washed several times with distilled water and then hydrolyzed in a 0.1% cellulase solution at 40°C. The degree of recovery of the cells after digestion was 80~95%. The isolated endosperm cells were diluted to 2~10 ml according to the developmental stage of the endosperm, from which 5 subsamples (20 ml per subsample) were transferred to a counting chamber (1-cm^2^ area). The endosperm cell number in 10 grids per counting chamber was counted using a light microscope. The number of nuclei was counted as the number of endosperm cells. The total endosperm cell numbers were calculated using the following equation (Liang *et al.*, 2001):

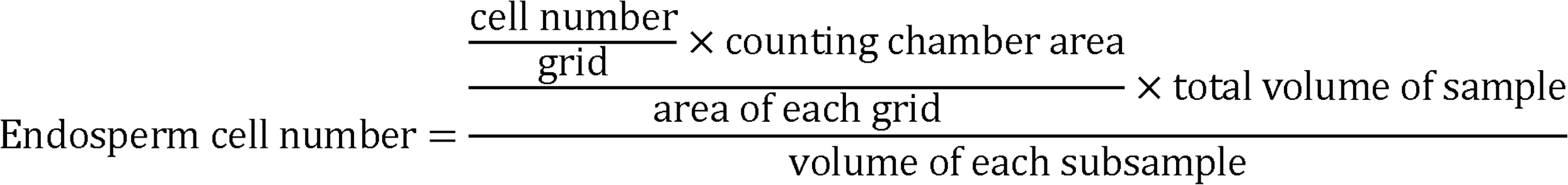

### Gene expression analysis

Total RNA was extracted from the grains (10~20 grains from five labeled panicles) using the RNAprep Pure Plant Kit (cat. no. DP441; Tiangen, Beijing, China). The HiScript II Q Select RT SuperMix (Vazyme, Nanjing, China) was used for cDNA synthesis. The transcript level of each gene was measured by qRT-PCR using the 7500 Real-Time PCR System (ABI) with the PowerUp™ SYBR® Green Master Mix (Thermo Fisher Scientific, San Jose, USA). Gene expression was quantified during the logarithmic phase using expression of the housekeeping gene *Ubq* (LOC_Os03g13170) as an internal control. Three biological replicates were performed for each experiment. The primer sequences used for qRT-PCR are described by Wang *et al.* (2013).

### Hormone quantification

The IAA and ABA levels were determined by Zoonbio Biotechnology Co., Ltd (Nanjing, China). Approximately 0.5 g of the samples were ground in a precooled mortar that contained 5 ml of extraction buffer composed of isopropanol/hydrochloric acid. The extract was shaken at 4°C for 30 min. Then, 10 ml of dichloromethane was added, and the sample was shaken at 4°C for 30 min and centrifuged at 13,000 rpm for 5 min at the same temperature. We extracted the lower organic phase. The organic phase was dried under N_2_, dissolved in 150 μl of methanol (0.1% methane acid) and filtered with a 0.22-μm filter membrane. The purified product was subjected to high-performance liquid chromatography-tandem mass spectrometry (HPLC-MS/MS) analysis. The HPLC analysis was performed using a ZORBAX SB-C18 (Agilent Technologies) column (2.1 mm × 150 mm; 3.5 mm). The mobile phase A solvent consisted of methanol/0.1% methanoic acid, and the mobile phase B solvent consisted of ultrapure water/0.1% methanoic acid. The injection volume was 2 μl. The MS conditions were as follows: the spray voltage was 4500 V; the pressure of the air curtain, nebulizer and aux gas were 15, 65 and 70 psi, respectively; and the atomizing temperature was 400°C.

### Enzyme activity assays

The dehulled grains (10 grains from five labeled panicles) were homogenized with a prechilled mortar and pestle in 100 mM HEPES buffer (pH 7.5) containing 8 mM MgCl_2_, 2 mM EDTA, 50 mM 2-mercaptoethanol, 12% (v/v) glycerol and 1% (w/v) polyvinylpyrrolidone (PVP). After centrifugation at 30,000 × g for 10 min at 4°C, the supernatant was desalted through a dialytic membrane. The dialysis buffer contained 5 mM HEPES-NaOH, pH 7.4, 5 mM MgCl_2_, 1 mM EDTA and 0.5 mM DTT. The enzyme activity of SUS (in the cleavage direction), SS and BE was determined as described by Nakamura *et al.* (1989), Jiang *et al.* (2003) and Tang *et al.* (2009). The grain starch contents were determined according to Lü *et al.* (2008).

### Protein blotting analysis

Rice grains (~10 grains from five labeled panicles of different plants) were homogenized in TEDM buffer (20 mM Tris/HCl, pH 7.5, 1 mM DTT, 5 mM EDTA and 10 mM MgCl_2_) containing a complete protease inhibitor cocktail (Roche). The homogenate was centrifuged at 6,000 × g for 30 min at 4°C to remove cellular debris, and the supernatant was clarified by centrifugation at 5,000 × g for 90 min at 4°C. The soluble proteins were separated by SDS—PAGE on a 10% gel and blotted onto a Polyvinylidene difluoride (PVDF) membrane (Millipore, Bedford, MA, USA). The DEP1/qPE9-1 (LOC_Os09g26999) and RGB1 (LOC_Os03g46650) antibodies were prepared by ABclonal (Wuhan, China). Serum was collected from a rabbit after multiple injections of the RGB1 (full-length) protein and a polypeptide (MEAPRPKSPPRYPDLC) of qPE9-1 as the antigen. The evaluation of antibody valence is shown in Figure S1.

### Statistical analysis

The data are presented as the mean ± SD. The SPSS 16.0 software was used for all statistical analyses. Statistical significance was determined for independent biological samples using Student’s t-test for comparison of two groups and one-way ANOVA for comparison of three or more groups. Differences were considered statistically significant when *P* < 0.05. An asterisk (*) is presented when *P* ◻ 0.05.

## Results

### *qPE9-1* positively controls grain size

Using QTL analysis, two independent research groups originally identified rice *qPE9-1* as controlling the panicle morphology and grain number per panicle (Huang *et al.*, 2009; Zhou *et al*, 2009). However, the biochemical and molecular mechanisms underlying control of the rice grain size and weight are largely unknown. In the present study, we compared the differences in grain size of Wuyunjing 8, which contains the *qpe9-1* allele (hereafter *qpe9-1*), and the *qPE9-1* transgenic line (hereafter *qPE9-1*). The results showed that the *qpe9-1* grain was approximately 12% shorter than that of *qPE9-1* (Fig. 1a and f) and that *qPE9-1* had a larger grain size during the grain-filling stage than *qpe9-1* (Fig. 1c). However, the width and thickness of the *qpe9-1* and *qPE9-1* grains did not differ significantly (Fig. 1b and f). The 1000-grain weight of *qPE9-1* was remarkable higher than that of *qpe9-1* (Fig. 1g). The mRNAs of *qPE9-1* are still present in *qpe9-1*, although neither the full-length protein (qPE9-1, predicted molecular weight 45.2 kDa) nor a truncated protein (qpe9-1, predicted molecular weight 21.5 kDa) was detected by immunoblotting with the anti-qPE9-1 antibody (Figs. 1d and e). These results indicate that the full-length qPE9-1 protein promotes the grain length in rice.

**Figure 1.**
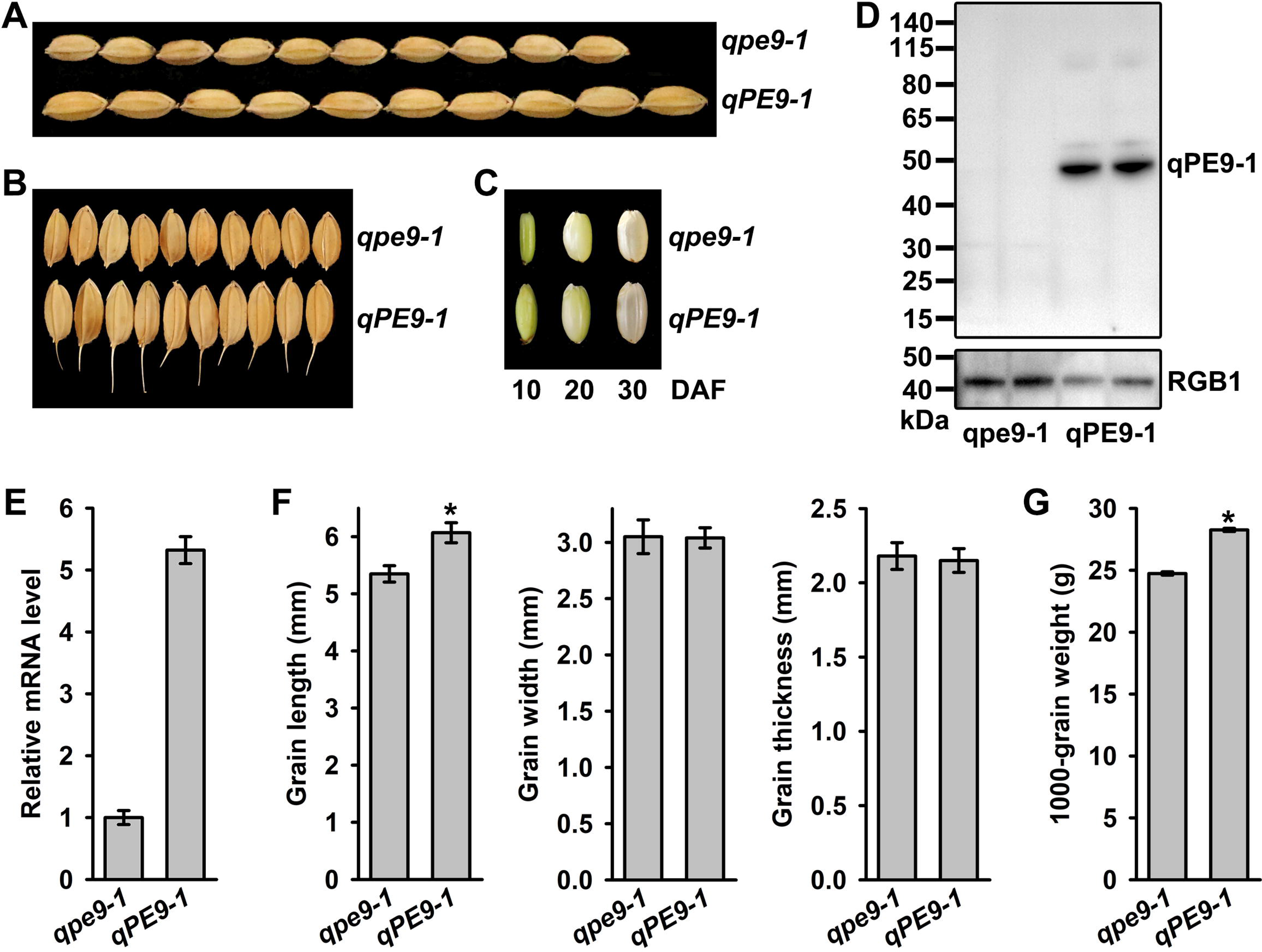
Grain performance of *qpe9-1* and *qPE9-1*. **a** Grain length; **b** Grain width; **c** Brown rice during grain development; **d** qPE9-1 protein expression in grains at 5 DAF was analyzed by incubating isolated proteins with polyclonal antibodies against qPE9-1 or RGB1 (as a loading control); **e** *qPE9-1* gene expression was monitored in grains at 5 DAF using qRT-PCR (n=3); **f** Comparisons between *qpe9-1* and *qPE9-1* with respect to the grain length, width and thickness; **g** The 1000-grain weight (n=3). DAF, days after flowering. Data are presented as the mean ± SD. “*” represents a significant difference at *P*<0.05.

### *qPE9-1* endosperm cell proliferation

Organ size and shape are determined by cell proliferation and expansion (Orozco-Arroyo *et al.*, 2015). We compared the numbers of endosperm cells in cross-sections of mature grains from *qpe9-1* and *qPE9-1* (Fig. 2a). No significant difference in the endosperm cell area was observed between the *qpe9-1* and *qPE9-1* lines (Fig. 2b). We also found that the number of endosperm cells per grain increased rapidly during early grain filling and that the peak values were reached nine days after flowering (9 DAF). The number of endosperm cells per grain of *qPE9-1* was significantly higher than that of *qpe9-1* (Fig. 2c). The long and short axes of the *qpe9-1* endosperm cells were slightly shorter than those of *qPE9-1*, but the differences were not significant (Fig. 2d). There findings suggest that *qPE9-1* has no obvious effect on cell proliferation during grain growth and development and that the increase in endosperm (grain) size in *qPE9-1* results mainly from an increase in cell numbers during the early stage of grain development.

**Figure 2.**
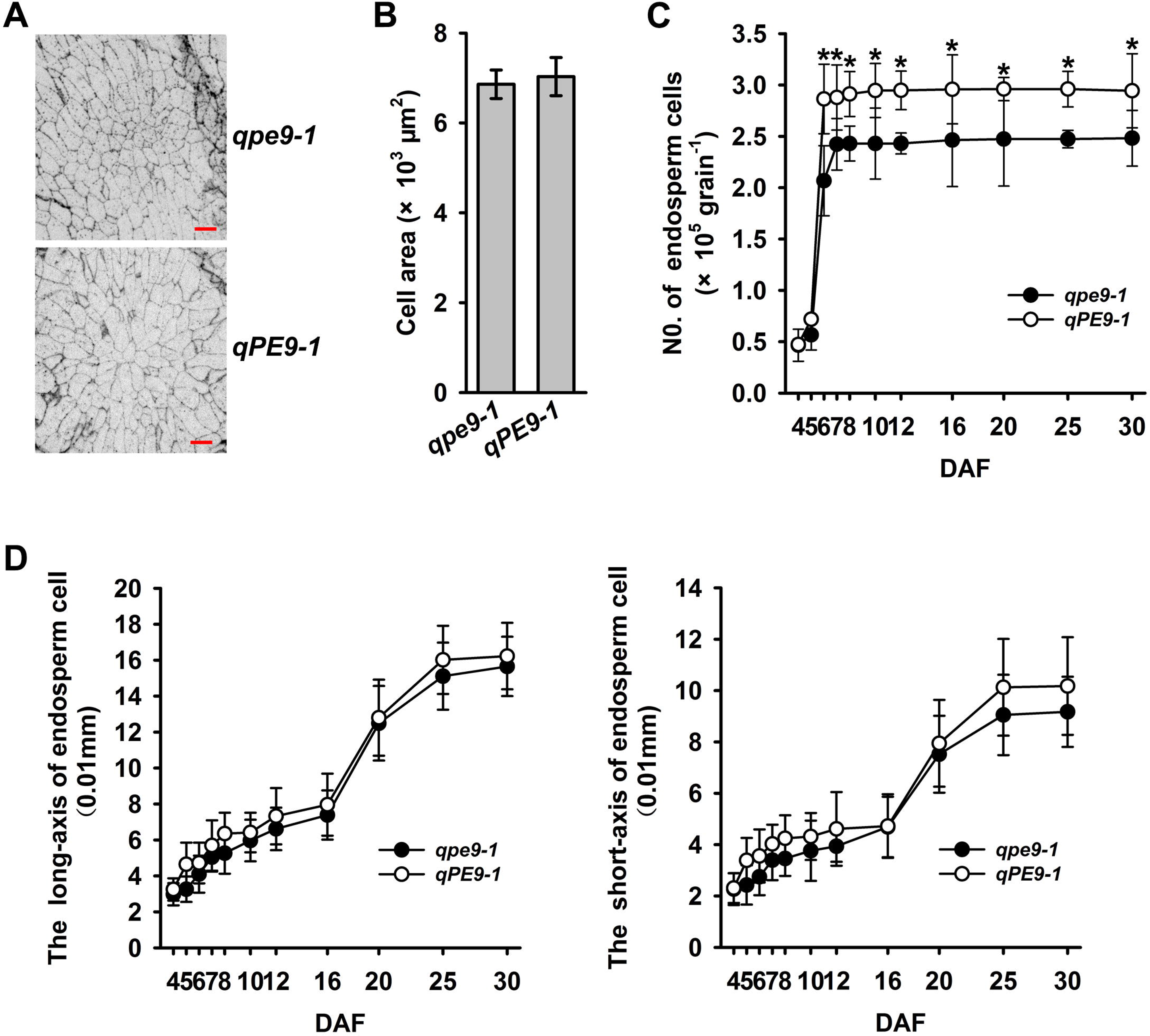
Histological analyses of endosperms at maturity and changes in endosperm size after fertilization in *qpe9-1* and *qPE9-1*. **a** Cross-sections of the endosperm between the dorsal and central points showing the cell sizes and numbers. Scale bars, 100 μm; **b** Comparison of cell numbers in the endosperm cross-sections (n=5 endosperms from 5 panicles); **c** Changes in the numbers of endosperm cells during grain filling (n=10 endosperms from 5 panicles); **d** The long and short axes of endosperm cells during grain development. Data are presented as the mean ± SD. “*” represents a significant difference at *P* < 0.05.

### *qPE9-1* enhances starch biosynthesis in rice grains

The starch content of the grains increased rapidly after flowering in both genotypes. However, the grain of the *qPE9-1* line contained a much higher starch content than that of the *qpe9-1* line, and the final starch content measured in the *qPE9-1* grain was approximately 25% higher than that in *qpe9-1* (Fig. 3a). Interestingly, the maximum starch content was measured at 27 DAF in *qpe9-1* and remained relatively stable until the end of grain filling, whereas the starch content in *qPE9-1* increased continuously until 33 DAF (Fig. 3a). The rate of starch accumulation increased rapidly after flowering, reached its maximal level at 15 DAF and then decreased as the grain-filling process continued. Again, the rate of starch accumulation during grain filling was almost always higher in *qPE9-1* than in *qpe9-1*, especially from the mid to late grain-filling stage (from 18 to 33 DAF, Fig. 3b). These results clearly indicate that *qPE9-1* positively controls the starch accumulation process during grain filling and prolongs the duration of the grain-filling process. Furthermore, the total starch content in *qPE9-1* flour was significantly higher than that of *qpe9-1*, suggesting that *qPE9-1* increased conversion of sucrose to starch.

**Figure 3.**
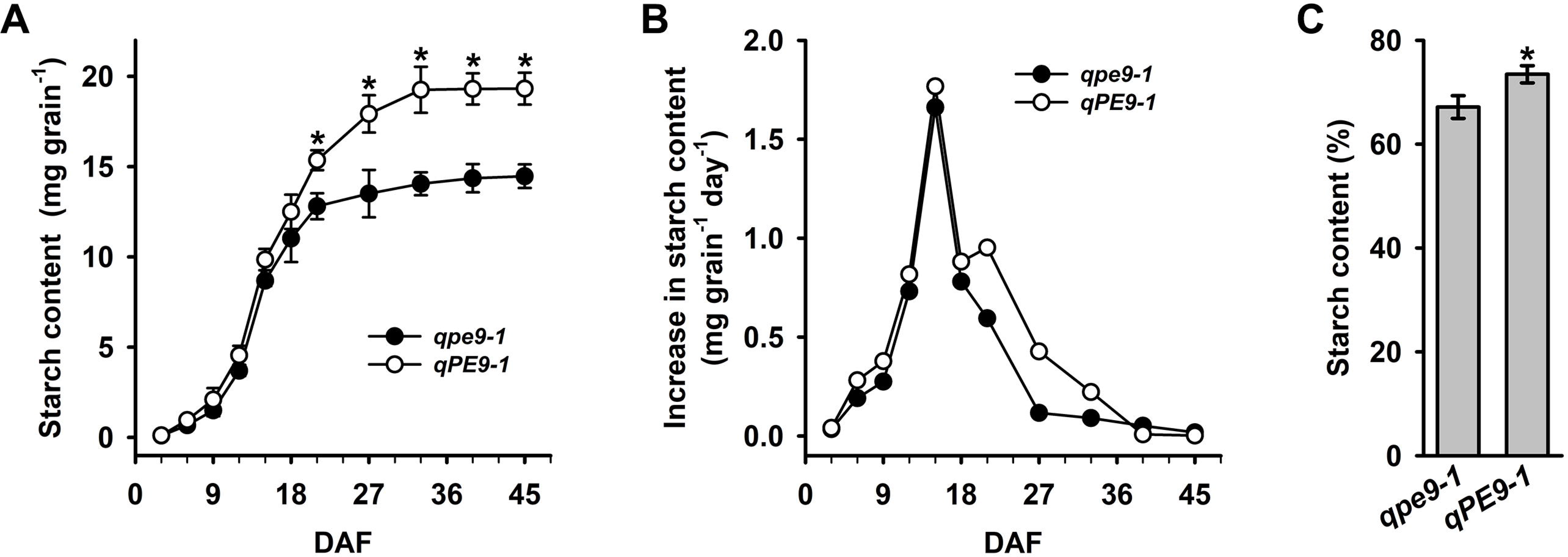
Starch accumulation during grain filling. **a** Starch accumulation of grains during the grain-filling stages (n=5); **b** The rate of starch accumulation during grain filling; **c** The total starch content in the flour (n=5). Data are presented as the mean ± SD. “*” represents a significant difference at *P* < 0.05.

### *qPE9-1* stimulates the expression of genes encoding grain-filling-related enzymes

Sucrose synthase (SUS), invertase (INV), ADP-glucose pyrophosphorylase (AGP), starch synthases (SS and GBSS), branching enzyme (BE) and debranching enzyme (DBE) are well-known to play key roles in the regulation of sucrose degradation, ADP-glucose and starch biosynthesis (Liang *et al.*, 2001; Lü *et al.*, 2008; Tang *et al.*, 2009; also see reviews by Tetlow *et al.*, 2004; Keeling & Myers, 2010; Zeeman *et al.*, 2010). In this study, we found that the *OsSUS3*, *OsSSIIa* and *OsBEIIb* transcript levels were significantly higher in the *qPE9-1* grains than in the *qpe9-1* grains. Then, we examined the expression of these genes during the entire grain-filling stage. The results showed that the transcript levels of these genes were significantly higher in the *qPE9-1* grains than in the *qpe9-1* grains, especially at the later grain-filling stage (Fig. 4a). These results suggest that the molecular mechanisms that regulate sucrose degradation and starch biosynthesis may differ between *qpe9-1* and *qPE9-1*, which may explain the differences in the final grain weight between the two genotypes.

**Figure 4.**
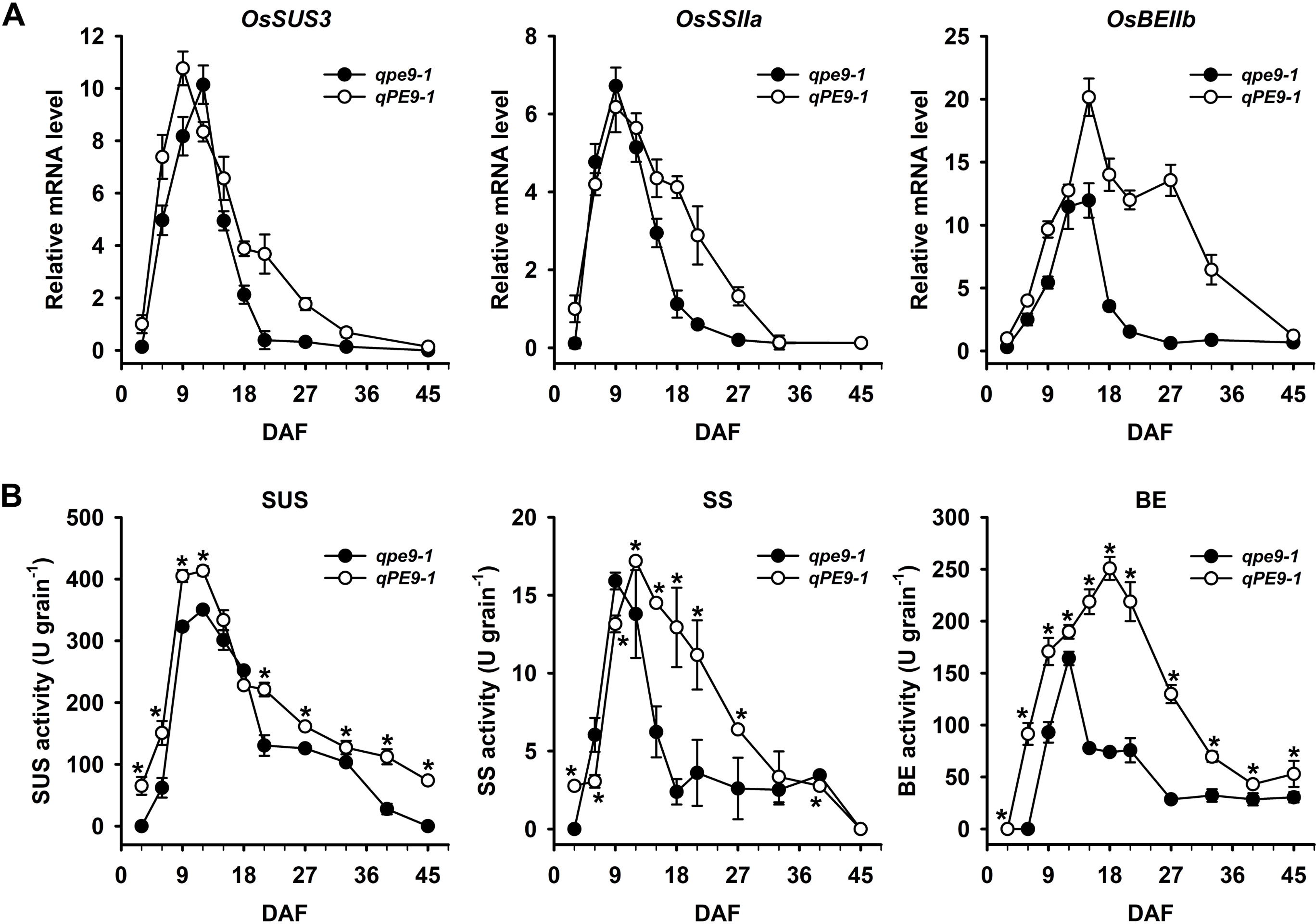
Expression of several starch biosynthesis genes and changes in the activities of these enzymes during grain filling. **a** *OsSUS3*, *OsSSIIa* and *OsBEIIb* expression levels during grain filling (n=3); **b** Changes in the SUS, SS and BE activities during grain filling (n=3), 1 U= 1 μg/min/mg protein. Data are presented as the mean ± SD. “*” represents a significant difference at *P* < 0.05.

To elucidate the roles of these genes at the biochemical level, the activities of several enzymes encoded by these genes were measured in the grains of the two genotypes during grain filling. As shown in Fig. 4b, significant differences in SUS, SS, GBSS and BE activity were observed between *qpe9-1* and *qPE9-1*; the dynamic changes in the activities of these enzymes during grain filling were very similar to the expression patterns of the encoded genes. Taken together, our results suggest that *qPE9-1* plays key roles in controlling grain filling through its effects on gene expression and the activities of the encoded enzymes related to sucrose metabolism and starch biosynthesis during grain filling.

### *qPE9-1* enhances the accumulation of ABA and IAA in the endosperm during grain filling

Rice grain filling is a complicated process that involves the interconversion of many metabolites and the metabolism of bioactive compounds, including plant hormones. The rice grain-filling process is highly regulated by plant hormones (Tang *et al.*, 2009; Zhu *et al.*, 2011; Zhang *et al*, 2016). We were interested in investigating whether *qPE9-1* controlled the grain-filling process through its effects on plant hormone homeostasis. Therefore, in the present study, the contents of two plant hormones were measured during grain filling. Significant changes in the endogenous ABA and IAA contents in the grains were found during grain filling (Figs. 5a and c). The ABA content increased during the early grain filling stage until 12 DAF and then decreased until the end of grain filling. The endogenous ABA content in the *qPE9-1* grains was always higher than that in *qpe9-1* (Fig. 5a). Very similar patterns of dynamic changes in the IAA content were observed, with the *qPE9-1* grains having higher IAA levels than *qpe9-1* during the grain-filling stages (Fig. 5c). The transcript levels of rice *OsNCED5*, which encodes a key enzyme catalyzing ABA biosynthesis [9-cis-epoxycarotenoid dioxygenase (*NCED*)], during grain filling were higher in *qPE9-1* than in *qpe9-1* from 3 to 15 DAF (Fig. 5b). The change in the transcript level of one important IAA biosynthesis gene (*OsTAR1*) was highly synchronized with changes in the grain IAA level (Fig. 5d). These results suggest that ABA and IAA may be involved in grain-filling regulation and that *qPE9-1* may exert its effects by changing the biosynthesis of these hormones during grain filling.

**Figure 5.**
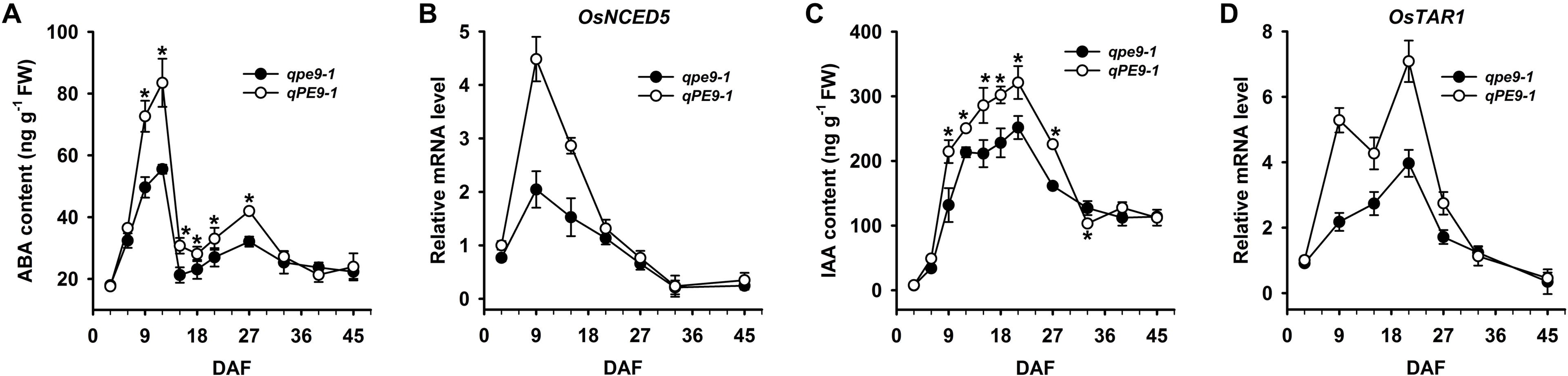
Changes in the ABA and IAA contents and the expression of genes encoding key enzymes catalyzing their biosynthesis during grain filling. **a** Changes in the ABA level in rice endosperm cells during seed development (n=3); **b** Changes in the expression profile of the ABA synthesis gene *OsNCED5* in rice endosperm cells during seed development (n=3); **c** Changes in the IAA levels in rice endosperm cells during seed development (n=3); **d** Changes in the expression profile of the IAA synthesis gene *OsTAR1* in rice endosperm cells during seed development (n=3). Data are presented as the mean ± SD. “*” represents a significant difference at *P* < 0.05.

### *qPE9-1* regulates starch biosynthesis in rice grains by ABA and IAA

To verify the relationships between *qPE9-1* and the plant hormones ABA and IAA during the grain-filling stages, exogenous ABA and IAA were applied to the developing grains during the filling stage, and starch accumulation and gene expression were analyzed. As shown in Fig. 6, all treatments had stimulatory effects on *OsSUS3*, *OsBEIIb* and *OsSSIIa* gene expression and the activities of their encoded enzymes in the *qPE9-1* grains, but ABA application had no effects on starch accumulation or gene expression in the *qpe9-1* grains (i.e., ABA had either inhibitory or no effects on starch accumulation). Compared with *qpe9-1* grains, *qPE9-1* grains showed enhanced starch accumulation upon treatment with either exogenous ABA or IAA, and this stimulatory effect of IAA was much more significant than that of ABA (Fig. 6a). Application of IAA and ABA also stimulated *OsSUS3*, *OsBEIIb* and *OsSSIIa* gene expression and the activities of their encoded enzymes in *qPE9-1* (Figs. 6b and c). These results indicate that starch accumulation in *qPE9-1* grains is largely, if not completely, dependent on high ABA and IAA levels.

**Figure 6.**
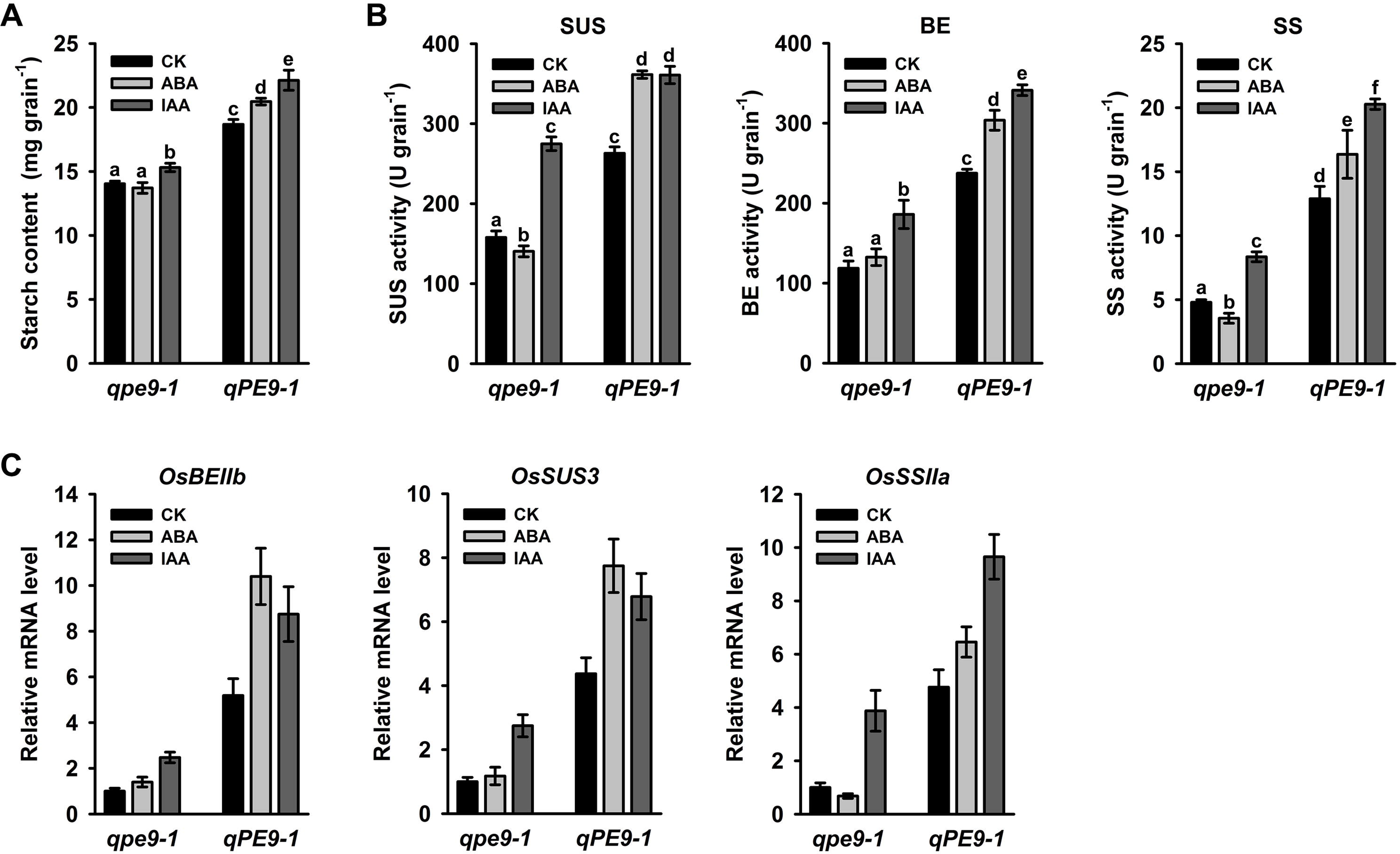
Effects of exogenous ABA and IAA applications on starch biosynthesis. **a** Effects of exogenous ABA and IAA on starch accumulation (n=5); **b** Effects of exogenous ABA and IAA on the SUS, SS and BE activities at 21 DAF (n=3), 1 U= 1 μg/min/mg protein; **c** Effects of exogenous ABA and IAA on *OsSUS3*, *OsSSIIa* and *OsBEIIb* expression at 21 DAF (n=3). Data are presented as the mean ± SD. “*” represents a significant difference at *P* < 0.05.

## Discussion

The rice *DEP1/qPE9-1* gene was formerly identified by two independent research groups as a gene that controls panicle erectness, grain number per panicle and, consequently, grain yield (Huang *et al.*, 2009; Zhou *et al.*, 2009). Sequential alignment at the amino acid level showed that *qPE9-1* was a homolog of *AGG3*, which is a γ subunit of the *Arabidopsis* heterotrimeric G protein, although its similarity and degree of homology were not as high. AGG3 is a recently discovered novel protein that is almost twice as large as a typical G_γ_; protein. This protein exhibits a high degree of similarity with canonical G_γ_ proteins at the N-terminus, whereas the C-terminus is plant-specific and contains an extremely high number of cysteine residues (Chakravorty *et al.*, 2011; Botella, 2012). In plants, the G_γ_ and G_β_ subunits are tightly bound physically and exert their effects as a dimer in both the active and inactive states of the heterotrimeric G protein. Recent studies showed that based on the interaction between DEP1/qPE9-1 and RGB1 (G_β_ subunit of rice), DEP1/qPE9-1 is a G_γ_ protein in rice (Botella, 2012; Sun *et al.*, 2014).

The functions and molecular mechanisms of DEP1/qPE9-1 as a new G_γ_ subunit in rice are largely unknown. Because *DEP1/qPE9-1* has been cloned and identified as a gene controlling the panicle size and rice shape (Huang *et al.*, 2009; Zhou *et al.*, 2009), we can reasonably assume that *DEP1/qPE9-1* may play important roles in regulating grain development and the grain-filling process. Rice grain formation and grain filling are rather complicated processes that involve approximately 21,000 genes, including 269 genes that are closely related to various physiological and biochemical pathways (Zhu *et al.*, 2003). Proteomic research indicates that 123 proteins and 43 phosphoproteins are involved in regulation of the grain-filling process (Zhang *et al.*, 2014). Furthermore, the grain-filling process is also regulated by plant hormones (e.g., ABA, IAA, cytokinin and ethylene), which fluctuate considerably during the grain-filling period (Yang *et al.*, 2001; 2003; 2006; Tang *et al.*, 2009). Although the carbohydrate interconversion biochemical pathway has clearly been elucidated during grain filling, the biochemical and molecular mechanisms controlling rice grain size and weight are largely unknown.

Overexpression of *Arabidopsis* atypical G_γ_ (AGG3) promotes seed and organ growth by increasing cell proliferation, and AGG3-mutated lines have a small seed size (Chakravorty *et al.*, 2011; Li *et al.*, 2012; Roy Choudhury *et al.*, 2013). Similar results have been reported for rice plants. *GRAIN SIZE 3 (GS3*) and *DENSE AND ERECT PANICLE1* (*DEP1*)/*PANICLE ERECTNESS* (*qPE9-1*) are involved in controlling either the grain size or panicle size in rice (Fan *et al.*, 2006; Huang *et al.*, 2009; Zhou *et al.*, 2009). However, in contrast to *Arabidopsis, GS3* negatively regulates the grain size, whereas *DEP1/qPE9-1* positively regulates both the grain and panicle size (Fan *et al.*, 2006; Huang *et al.*, 2009; Zhou *et al.*, 2009). Why this considerable difference exists between these plant species remains unclear and should be studied and elucidated in the future.

A comparison of the panicle morphology, grain size and final grain weight clearly indicated that *qpe9-1* had a smaller size and lower grain weight than *qPE9-1*, implying that *qPE9-1* positively regulated the grain size and final grain weight (Fig. 1). Our previous research and that conducted by other groups revealed that the grain size and final grain weight were largely determined by two factors/processes: the number and size of endosperm cells (i.e., the sink size) and the grain-filling process, including the grain-filling rate and duration (i.e., the sink activity; Liang *et al.*, 2001). The smaller grain size and lower final grain weight of *qpe9-1* are a result of fewer endosperm cells and lower starch accumulation after flowering, the latter is partly due to reducing the duration of the grain-filling process (Figs. 2 and 3). However, the mechanisms underlying how *qPE9-1* controls endosperm cell proliferation and starch accumulation are unknown.

More than ten proteins/enzymes are directly involved in the biochemical pathways of carbohydrate interconversion and starch biosynthesis during rice grain filling (Tetlow *et al.*, 2004; Ohdan *et al.*, 2005; Zhu *et al.*, 2011). What is the underlying cause of lower starch accumulation in *qpe9-1* aside from the lower numbers of endosperm cells? In other words, do significant differences exist in the expression of key protein/enzyme-encoding genes or the activity of these key enzymes between the two genotypes? To answer these questions, we compared differences in the expression of genes encoding several key enzymes that catalyzed carbohydrate interconversion and starch biosynthesis between the two genotypes used in this experiment. Our results clearly indicate that the lower expression levels of several key genes that encode enzymes catalyzing starch biosynthesis and the lower activities of these enzymes in the *qpe9-1* grains during the late grain filling stage are the most important factors resulting in the lower grain weight (Fig. 4).

The grain-filling process is a highly regulated process that involves both genetic and environmental factors. Plant hormones play important roles in grain growth and development (Tang *et al.*, 2009; Zhu *et al.*, 2011; Zhang *et al.*, 2016). However, little is known about the detailed mechanisms by which plant hormones regulate the grain-filling process. Yang *et al.* (2001) suggested that an altered hormonal balance, especially a decrease in GA and an increase in ABA, enhanced the remobilization of prestored carbon to the grains and accelerated the grain-filling rate. In addition, IAA treatment increased spikelet growth and development in distal branches (Patel & Mohapatra, 1992). Recently, studies in peas provided direct evidence that auxin was required for a normal seed size and starch accumulation. The mutant of *TAR2*, which is an IAA biosynthesis gene, induces the formation of small seeds with reduced starch content and a wrinkled phenotype at the dry stage. Application of the synthetic auxin 2,4-D partially reversed the wrinkled phenotype but did not restore the starch content of the mutant seeds to that of the WT (McAdam et al., 2017). Our results showed considerable differences in the dynamics of the endogenous hormone levels in the grains during grain filling, and the endogenous ABA and IAA levels were positively related to grain filling during the entire grain-filling stage (Fig. 6). The levels of these endogenous plant hormones were significantly lower in the *qpe9-1* than in the *qPE9-1* grains during the grain-filling stages (Figs. 5a and c). Based on the effects of exogenous ABA and IAA on starch accumulation and the expression of genes that encode several key enzymes catalyzing starch biosynthesis, we conclude that the positive control of *qPE9-1* on the grain-filling process occurs largely through the biosynthesis of endogenous plant hormones.

Rice grain filling is a very complicated process that involves photoassimilate (mainly sucrose) translocation from photosynthetic sources (i.e., leaves and leaf sheaths), sucrose degradation, transmembrane transport and starch synthesis in the grains (i.e., the sink, Liang *et al.*, 2001; Lü *et al.*, 2008; Tang *et al.*, 2009). Approximately twenty enzymes/proteins have been reported to be involved in these biochemical processes (Tetlow *et al.*, 2004; Ohdan *et al*, 2005; Zhu *et al.*, 2003). However, the mechanism of starch biosynthesis regulation in grains is not well understood. This research showed that the novel G protein γ subunit *qPE9-1* increased the numbers of endosperm cells and positively regulated starch biosynthesis, which enhanced the grain size and weight. *qPE9-1* also enhanced the accumulation of ABA and IAA. In turn, these hormones regulate the expression of genes encoding several key enzymes that catalyze starch biosynthesis during the late grain filling stage and consequently affect the final grain weight.

## Supplementary Data

**Figure S1.** The evaluation of antibody valence. **a** Analysis of the RGB1 protein using a RGB1 antibody; **b** Analysis of a peptide fragment of the DEP1/qPE9-1 protein using a DEP1/qPE9-1 antibody; **c** Detection of the His-tagged DEP1/qPE9-1 protein using both DEP1/qPE9-1 and His tag antibodies.

## Acknowledgements

We thank Dr. Cunxu Wei (Yangzhou University) for the fruitful discussions and valuable suggestions on this study.

## Reference

Botella JR. 2012. Can heterotrimeric G proteins help to feed the world? Trends in Plant Science 17, 563–568

Chakravorty D, Trusov Y, Zhang W, Acharya BR, Sheahan MB, McCurdy DW, Assmann SM, Botella JR. 2011. An atypical heterotrimeric G-protein γ-subunit is involved in guard cell K^+^-channel regulation and morphological development in Arabidopsis thaliana. Plant Journal 67, 840–851

Fan C, Xing Y, Mao H, Lu T, Han B, Xu C, Li X, Zhang Q. 2006. *GS3*, a major QTL for grain length and weight and minor QTL for grain width and thickness in rice, encodes a putative transmembrane protein. Theoretical and Applied Genetics 112, 1164–1171

Huang X, Qian Q, Liu Z, Sun H, He S, Luo D, Xia G, Chu C, Li J, Fu X. 2009. Natural variation at the *DEP1* locus enhances grain yield in rice. Nature Genetics 41, 494–497

Jiang D, Cao W, Dai T, Jing Q. 2003. Activities of key enzymes for starch synthesis in relation to growth of superior and inferior grains on winter wheat (*Triticum aestivum* L.) spike. Plant Growth Regulation 41, 247–257

Keeling PL, Myers AM. 2010. Biochemistry and genetics of starch synthesis. Annual Review of Food Science and Technology 1, 271–303

Li S, Liu Y, Zheng L, et al. 2012. The plant-specific G protein γ subunit AGG3 influences organ size and shape in *Arabidopsis thaliana*. New Phytologist 194, 690–703

Liang J, Zhang J, Cao X. 2001. Grain sink strength may be related to the poor grain filling of *indica-japonica* rice (*Oryza sativa*) hybrids. Physiologia Plantarum 112, 470–477

Lü B, Guo Z, Liang J. 2008. Effects of the activities of key enzymes involved in starch biosynthesis on the fine structure of amylopectin in developing rice (*Oryza sativa* L.) endosperms. Science in China. Series C 51, 863–871

McAdam EL, Meitzel T, Quittenden LJ, Davidson SE, Dalmais M, Bendahmane AI, et al. 2017. Evidence that auxin is required for normal seed size and starch synthesis in pea. New Phytologist 216, 193–204

Milligan G, Kostenis E. 2006. Heterotrimeric G-proteins: a short history. British Journal of Pharmacology 147 Suppl 1, S46–55

Nakamura Y, Yuki K, Park S-Y, Ohya T. 1989. Carbohydrate Metabolism in the Developing Endosperm of Rice Grains. Plant and Cell Physiology 30, 833–839

Ohdan T, Francisco PB, Sawada T, Hirose T, Terao T, Satoh H, Nakamura Y. 2005. Expression profiling of genes involved in starch synthesis in sink and source organs of rice. Journal of Experimental Botany 56, 3229–3244

Orozco-Arroyo G, Paolo D, Ezquer I, Colombo L. 2015. Networks controlling seed size in *Arabidopsis*. Plant Reproduction 28, 17–32

Patel R, Mohapatra PK. 1992. Regulation of spikelet development in rice by hormones. Journal of Experimental Botany 43, 257–262

Roy Choudhury S, Riesselman AJ, Pandey S. 2013. Constitutive or seed-specific overexpression of *Arabidopsis G-protein γ subunit 3* (*AGG3*) results in increased seed and oil production and improved stress tolerance in *Camelina sativa*. Plant Biotechnology Journal 12, 49–59

Singh BK, Jenner CF. 1982. A modified method for the determination of cell number in wheat endosperm. Plant Science Letters 26, 273–278

Sun H, Qian Q, Wu K, Luo J, Wang S, Zhang C, Ma Y, Liu Q, Huang X, Yuan Q, et al. 2014. Heterotrimeric G proteins regulate nitrogen-use efficiency in rice. Nature Genetics 46, 652–656

Tang T, Xie H, Wang Y, Lü B, Liang J. 2009. The effect of sucrose and abscisic acid interaction on sucrose synthase and its relationship to grain filling of rice (*Oryza sativa* L.). Journal of Experimental Botany 60, 2641–2652

Tetlow IJ, Morell MK, Emes MJ. 2004. Recent developments in understanding the regulation of starch metabolism in higher plants. Journal of Experimental Botany 55, 2131–2145

Urano D, Alan AM. 2014. Heterotrimeric G Protein-Coupled Signaling in Plants. Annual Review of Plant Biology 65, 365–84

Urano D, Chen J-G, Botella JR, Jones AM. 2013. Heterotrimeric G protein signalling in the plant kingdom. Open biology 3, 120186

Wang J-C, Xu H, Zhu Y, Liu Q-Q, Cai X-L. 2013. OsbZIP58, a basic leucine zipper transcription factor, regulates starch biosynthesis in rice endosperm. Journal of Experimental Botany 64, 3453–3466

Yang J, Zhang J, Wang Z, Zhu Q. 2003. Hormones in the grains in relation to sink strength and postanthesis development of spikelets in rice. Plant Growth Regulation 41, 185–195

Yang J, Zhang J, Wang Z, Liu K, Wang P. 2006. Post-anthesis development of inferior and superior spikelets in rice in relation to abscisic acid and ethylene. Journal of Experimental Botany 57, 149–160

Yang J, Zhang J, Wang Z, Zhu Q, Wang W. 2001. Hormonal changes in the grains of rice subjected to water stress during grain filling. Plant Physiology 127, 315–323

Zeeman SC, Kossmann J, Smith AM. 2010. Starch: its metabolism, evolution, and biotechnological modification in plants. Annual Review of Plant Biology 61, 209–234

Zhang W, Cao Z, Zhou Q, Chen J, Xu G, Gu J, Liu L, Wang Z, Yang J, Zhang H. 2016, Grain Filling Characteristics and Their Relations with Endogenous Hormones in Large- and Small-Grain Mutants of Rice. PLoS ONE 11, e0165321

Zhang Z, Zhao H, Tang J, Li Z, Li Z, Chen D, Lin W. 2014. A proteomic study on molecular mechanism of poor grain-filling of rice (*Oryza sativa* L.) inferior spikelets. PLoS ONE 9, e89140

Zhou Y, Zhu J, Li Z, Yi C, Liu J, Zhang H, Tang S, Gu M, Liang G. 2009. Deletion in a quantitative trait gene *qPE9-1* associated with panicle erectness improves plant architecture during rice domestication. Genetics 183, 315–324

Zhu G, Ye N, Yang J, Peng X, Zhang J. 2011. Regulation of expression of starch synthesis genes by ethylene and ABA in relation to the development of rice inferior and superior spikelets. Journal of Experimental Botany 62, 3907–3916

Zhu T, Budworth P, Chen W, Provart N, Chang H-S, Guimil S, Su W, Estes B, Zou G, Wang X. 2003. Transcriptional control of nutrient partitioning during rice grain filling. Plant Biotechnology Journal 1, 59–70

